# Evaluation of parasitic contamination and effects on estimates of mitochondrial enzymatic activities in infected fish livers

**DOI:** 10.1101/2025.02.25.640121

**Authors:** Tommy Pepin, Vincent Mélançon, Sandra Ann Binning, Sophie Breton

## Abstract

Parasites can impair host performance through various physiological processes, including alterations to host metabolism. Since mitochondria are responsible for cellular energy production, it is likely that disruptions in host cellular metabolism contribute to changes in metabolism at the organismal level. However, some studies investigating parasite-induced alteration in cellular metabolism have not taken the presence of parasites in the target tissues into account, potentially biasing results. It is therefore critical to confirm that the measured enzyme activities reflect those of the hosts rather than the parasites themselves. Here, we tested a parasite extraction protocol to evaluate the extent to which parasite contamination impacts estimates of cellular enzyme activities in hepatic tissues of wild pumpkinseed sunfish (*Lepomis gibbosus*) infected with bass tapeworm cestodes (*Proteocephalus ambloplitis*). We tested four treatments: uninfected livers, cleaned infected livers, infected livers (repopulated) and parasites alone. We then compared the activity of key metabolic enzymes among groups. PCR tests were used to assess parasitic contamination of samples after applying the parasite extraction protocol on hepatic tissue. Enzyme activities of cleaned livers and contaminated livers were similar despite PCR tests revealing contamination. The intensity of cestode infection also did not influence enzyme activity, which suggests that parasite presence in liver tissues does not impact the accuracy of the enzyme activity estimates. These results suggest that studying the organs of heavily parasitized individuals is possible. Nevertheless, we recommend that our cleaning protocol is applied to infected organs to avoid any perception of biases in highly infected individuals.

## Introduction

Parasitism is one of the most widespread lifestyles on Earth, accounting for nearly half of all described species (Poulin and Morand 2000; Dobson et al. 2008). Cestodes are common parasitic flatworms that mainly live in the gastrointestinal tract and abdominal cavity of a wide range of host species (Mackiewicz 1988; Scholz et al. 2021). In some fish species, cestodes infect the liver and can have a wide range of performance impacts, such as alterations to metabolic activities, swimming speed, responsiveness to certain stimuli and fecundity (Meakins and Walkey 1975; Blake et al. 2006; Heins et al. 2014; Msafiri et al. 2014; Guitard et al. 2022). However, the mechanisms underlying cestode-induced changes in host performance remain poorly understood. Parasite infections impose an energetic burden on their hosts by triggering immune responses, causing tissue damage, and/or reallocating energy reserves (Lemly and Esch 1984; Delahay et al. 1995; Marcogliese 2004). For example, in the three-spined stickleback (*Gasterosteus aculeateus*), oxygen consumption rate, a proxy of whole-organism metabolic rate, increases with the number of cestodes, *Scistocephalus solidus,* infecting their body cavity (Meakins and Walkey 1975; Tarallo et al. 2021). Conversely, in pumpkinseed sunfish (*Lepomis gibbosus*), both standard and maximum metabolic rate (SMR and MMR) decrease with increasing density of the cestode *Proteocephalus ambloplitis* (Guitard et al. 2022). Although the underlying mechanisms driving these changes in host performance remain unclear, it is likely that infection-induced alterations in host cellular metabolism play a significant role.

To understand how parasitism alters host cellular metabolism, it is crucial to study the mitochondria, given their central role in ATP production - the primary source of energy produced by cellular respiration. ATP production occurs via oxidative phosphorylation (OXPHOS) in the inner mitochondrial membrane (Babcock 1999; Osellame et al. 2012). Some studies have found links between parasite infections and host cellular metabolism. For instance, in pumpkinseed sunfish, helminth infections have been associated with variations in the cellular metabolism of different tissues (Mélançon et al. 2023; Levet et al. 2024). Sabbagh et al (2024) specifically examined enzymatic activity in livers infected with *P. ambloplitis* and found links between cestode density and altered cellular metabolism across populations from three lakes. The liver plays an important role in several major physiological processes, including glycogen storage, lipid metabolism and protein synthesis (Tarasenko and McGuire 2017) and is a common target for some parasites (Deslyper et al. 2019). Consequently, parasite infections may contribute to the altered metrics often observed in heavily infected hosts (Laidley et al. 1988; Shaw et al. 2008; Nadler et al. 2021; Sabbagh et al. 2024). However, measures of enzyme activity or other metrics taken directly from infected tissues may be subject to bias. Some fish livers tested in Sabbagh et al (2024) were heavily infected with cestodes (2 populations: median = 26 and 37 and max = 179 and 180), which complicated the retrieval of host liver tissues. It is thus possible that the enzyme activities measured reflected contributions from the parasites themselves, even if parasites were removed.

This potential contamination with parasite tissue could undermine the reliability of enzymatic activity estimates from livers, especially when infection intensity is high. Indeed, Mélançon et al. (2023) and Levet et al. (2024) refrained from sampling pumpkinseed liver tissues due to high levels of parasite contamination. This is unfortunate, as heavily damaged tissues are likely to show the most pronounced alterations in host cellular metabolism. Since these tissues are also the most valuable for studying these changes, the development and validation of protocols to purify infected tissues and assess potential biases introduced by parasite contamination in liver samples are crucial to accurately assessing the impacts of extracellular parasites on host cellular metabolism.

The objective of this study is to develop and validate a protocol for parasite extraction and to assess parasite contamination in host livers, in order to better understand the effects of cestodes on hepatic tissue metabolism of wild, naturally infected fish. As a model organism, we used the pumpkinseed sunfish (*Lepomis gibbosus*), a freshwater species that is widespread and native to Eastern North America. Sunfish are commonly used in physiological and behavioural studies due to their susceptibility to parasitic helminth infections, notably by the cestode *Proteocephalus ambloplitis* (Hunter and Hamilton 1941; Wilson et al. 1996). We collected pumpkinseeds from Laurentian lakes in Quebec, which differ in cestode parasite prevalence. These cestodes have complex lifecycles with copepods as first intermediate hosts. Pumpkinseeds are second intermediate hosts and are infected through infected copepods ingestion (Fischer and Freeman 1969; Wilson et al. 1996). In sunfish, the cestodes develop in plerocercoids, the cestode larvae, and migrate from the gut wall to the abdominal cavity to the liver, spleen and/or gonads (Fischer and Freeman 1969, 1973). The cestodes enter their definitive hosts, small and largemouth bass (*Micropterus dolomieu* and *M. nigricans)* when sunfish are predated (Fischer and Freeman 1969).

We developed a “decontamination method” by comparing the activities of key metabolic enzymes from pumpkinseed hepatic tissues with and without parasites and performed PCR tests to evaluate parasitic contamination. Since we were able to remove all visible parasites from fish livers, we predicted that there should not be signals of contamination even though the livers were initially infected, and that the cellular metabolism of cleaned and contaminated livers would differ.

## Methods

### Ethical Statement

*Lepomis gibbosus* were collected and cared for with approval from the Université de Montréal’s animal care committee and the ministère de l’Environnement, de la Lutte contre les Changements Climatiques et de la Faune et des Parcs (Comité de Déontologie de l’Expérimentation sur les Animaux; Permit number: 23-025, Collection permit number: 2023-05-24-2124-15-S-P).

### Fish sampling

Wild pumpkinseed sunfish (*Lepomis gibbosus*) were collected with baited minnow traps from Lake Triton (n=10) and Lake Cromwell (n=40) in June 5-9, 2024, at the Laurentian Biological Field Station (SBL) in Quebec, Canada (lat. 45,98861, long. - 74,00585). Fish were selected between 70,00 to 110,00 mm in total length and with the goal to minimize non-targeted parasite infection (i.e. presence of visible trematodes *Uvulifer ambloplitis* & *Apophallus spp*.). Collected fish were euthanized in a eugenol solution (clove-oil; 4mL/L of water) and immediately submerged in ice in coolers to preserve tissue integrity. Fish were then brought to the SBL laboratory facilities and frozen at −20℃ prior to their transport (on ice) to the Université de Montréal’s Complexe des sciences laboratory facilities for dissection, enzymatic assays and PCR analysis.

### Fish dissection and liver parasite extraction

Fish were thawed on ice and measured with a 0-150 mm (± 0,01 mm) digital caliper. The total wet weight of each fish (including parasite biomass) was measured using an electronic balance (MSE225S, Sartorius Weighing Company, IL, ± 0,00001 g). Then, the liver was extracted on ice to maintain enzymatic capacities of the tissue. During liver dissection, cestodes and liver tissues were separated using dissection tweezers, and cestodes were counted under a dissecting microscope. The livers were screened twice to make sure that all cestodes were removed. All instruments were thoroughly cleaned with dish soap, rinsed with distilled water, and dried between each dissection. Liver tissues were classified into three different groups: (i) fish from Lake Triton, which were uninfected with parasites (control, n=10), (ii) fish from Lake Cromwell for which parasites were removed from the liver, counted, and kept separate for enzymatic analysis(cleaned; n=13), and (iii) fish from lake Cromwell for which parasites were removed from the liver, counted, and reintroduced into the liver tissues for enzymatic analysis (repopulated; n=13). Four additional naturally uninfected fish from Lake Cromwell were also included for comparison of their enzyme activities with the control group. Liver tissues and parasites were kept on ice in 5 mL cryogenic tubes, weighted using an electronic balance and stored at −80°C for further analyses. Fish were also screened for other parasites. After complete cestode removal, fish were re-weighted, and the liver mass was added to obtain the corrected fish weight (wet-weight without parasites) (Timi and Poulin 2020). Lastly, ten additional “cleaned” livers from fish from Lake Cromwell were frozen at −80°C (∼50 mg for each liver) for DNA extractions and PCR analysis to evaluate the possibility of parasite DNA contamination.

### Fish liver and parasite enzymatic activities

All samples were homogenized using a polytron (PT1200 E) as in Mélançon et al. (2023) and Sabbagh et al. (2024). Before being homogenized, parasites were washed with 1 ml of homogenization buffer (100 mM potassium phosphate [KP], 20 mM ethylenediamine tetraacetic acid (EDTA), pH 8,0) to remove any liver residual. Samples were vortexed for 10 seconds and then centrifuged at 3000 rpm at 4℃ for 10 minutes. The supernatant was carefully removed avoiding taking out any parasitic tissues. Afterwards, parasite samples were diluted 20 times their wet weight in the same homogenization buffer (100 mM KP, 20 mM EDTA, pH 8,0) as in Mélançon et al. (2023).

All liver samples were diluted 20 times their wet weight with the homogenization buffer. Enzymatic assays were performed on livers and parasites. The enzymes tested were the same as in Mélançon et al. (2023) and Sabbagh et al. (2024): cytochrome c oxidase (CCO) for oxidative phosphorylation (OXPHOS), citrate synthase (CS) for tricarboxylic cycle (TCA), lactate dehydrogenase (LDH) for anaerobic pathway and carnitine palmitoyltransferase (CPT) for fatty acid oxidation. For parasites, the quantity of tissue available restricted the enzymatic assays to CPT, CS and LDH. All enzymatic assays were run in duplicate and performed in 96-well plates using a microplate reader (Mithras LB 940, Berthold Technologies, Bad Wildbad, Germany) at room temperature (25℃).

### Cytochrome c oxidase (CCO) (Enzyme commission [EC] 7.1.1.9)

CCO is one of the complexes of the mitochondrial inner membrane (complex IV) and was chosen because of its implication in OXPHOS. CCO activity was measured at 550 nm over 5 min following the oxidation of cytochrome c (ɛ = 18,5 mM^-1^ cm^-^1). This was done using a 100 mM KP buffer containing 0.05 mM oxidized cytochrome c from equine heart, 4.5 mM sodium dithionate, and 0.03% Triton X-100, pH 8.0. Control reactions were performed by adding 10mM sodium azide to the sample and background activities were performed without sodium dithionate and then deducted to the activity of the assay (Thibault et al. 1997; Hunter-Manseau et al. 2019).

### Citrate synthase (CS) (EC 2.3.3.1)

CS is implied in the TCA cycle and was used as a quantitative enzyme marker of intact mitochondria content and indicator of the aerobic capacity. CS activity was measured at 412 nm following the conversion of 5,5’ dithiobis-2-nitrobenzoic acid (DTNB) into 2-nitro-5-thiobenzoic acid (TNB; ɛ = 14,15 mM^-1^ cm^-1^) over 6 min. The reaction solution consisted of 100 mM imidazole-HCl buffer with 0.1 mM DTNB, 0.1 mM acetyl-CoA, and 0.15 mM oxaloacetate, pH 8.0. A background without oxaloacetate was first performed followed by the final assay (Smith et al. 1985; Hunter-Manseau et al. 2019; Mélançon et al. 2023).

### Lactate dehydrogenase (LDH) (EC 1.1.1.27)

LDH catalyzes the reversible reaction of pyruvate into lactate, and is an important enzyme involved in the anaerobic metabolism (Khan et al. 2020). LDH activity was assessed at 340 nm by measuring the rate of NADH oxidation (ε = 6.22 mM^-1^ · cm^-1^) over 7 minutes using a KP buffer (100 mM) with NADH (0,168 mM) and pyruvate (0,4 mM), pH 7.0. Control reactions were performed by omitting the sample from the buffer (Thibault et al. 1997).

### Carnitine palmitoyltransferase (CPT) (EC 2.3.1.21)

CPT is involved in the transport of fatty acids across the inner mitochondrial membrane, allowing the oxidation of fatty acids into the mitochondrial matrix (Jogl et al. 2004). We used CPT activity as a control because parasites do not have the active form of the enzyme responsible for the beta-oxidation of fatty acids (Barrett and Körting 1977). We predicted that CPT activity should decrease with parasite load. CPT activity was assessed at 412 nm over 7 minutes by measuring the conversion of DTNB into TNB (ε = 14.15 mM-1・cm-1) using a 75 mM tris-HCl buffer, 1,5 mM EDTA with 0,25 mM DTNB, 0,035 mM PalmitoylCoA and 2 mM L-carnitine, pH 8.0. Control reactions were performed by omitting the sample (Thibault et al. 1997).

Enzymatic activities were normalized by organ wet weight or protein content for both fish hepatic tissues and parasites, and because normalization did not change the observed trends, total protein content was used to interpret the results. Total protein content (mg ML^-1^) was measured with the bicinchoninic acid method (Smith et al. 1985). Enzyme activities were expressed as U mg protein^-1^ where U represents 1 μmol of substrate transformed into product in 1 min.

### DNA extractions and PCR protocol

DNA extraction from liver tissues (n = 10; ∼50 mg each) was performed with the QIAGEN Blood & Tissue kit following the animal tissue protocol. DNA quality and quantity were examined via electrophoresis on a 1% agarose gel and a Biodrop μLITE spectrophotometer, respectively. The mitochondrial 16S gene was PCR-amplified in a total volume of 25 µL containing 2,5 µL of reaction buffer (10x), 0,5 µL dNTP, 1 µL of each primer (10 µM) specific to the *Proteocephalus ambloplitis* 16S gene (5’-AGGGCACAGAATCCTCCTTT-3’ and 5’-AGACTAGACTACTGCGCCAA-3’), 0,125 µL of Taq (5U/µL), 1 µL of DNA and 18,875 µL of ddH_2_O. PCR amplifications were performed with a TProfessional Basic Thermocycler in those conditions: a denaturation temperature of 95℃ for 2 min, then 35 cycles of 95℃ for 20 seconds, 55℃ for 40 seconds and 72℃ for 25 seconds ending with an elongation period of 5 min at 72℃. Two DNA samples from only parasites were used as a positive control to make sure the primers worked correctly, and sterile water was used as a negative control. PCR products were observed on a 1% agarose gel under UV light with SYBR green dye (Life Technologies).

### Statistical analysis

Linear models were used to compare enzymatic activities of the four naturally uninfected fish from Lake Cromwell and fish from Lake Triton using a Student t-test to determine whether uninfected fish have similar enzymatic activities regardless of their lake of origin. To evaluate the effect of parasite contamination on measures of enzymatic activities, LMs with enzymatic activities as the response variables and treatment (i.e control, cleaned, repopulated and parasite) as a fixed factor were used. When linear regression conditions were not respected, non-parametric Kruskall-Wallis tests by rank were accompanied by Wilcoxon-Mann-Whitney tests with Holm’s correction to compare pairwise rankings. When linear regression conditions were not respected, Student T-tests were replaced by non-parametric Wilcoxon-Mann-Whitney tests by rank. To determine whether enzymatic activities were affected by the density of worms present in a liver, we used a linear regression with a two-way interaction between liver cestode density and treatment (i.e. cleaned and repopulated) for all enzyme activities. Statistical analyses were performed with the version 4.2.3 of R (R Development Core Team) with ggplot2 package and any p-value < 0.05 were considered significant.

## Results

Fish from both lakes had a mean total length of 93,36 mm (± SE; ± 2,0685 mm). Fish from Lake Triton were not infected by cestodes and had a mean wet-weight of 19,47g (min-max ± SE; 10.30-32.48 ± 1,78g). Fish from Lake Cromwell had an 87% infection prevalence with abundance ranging from 0 to 54 (mean ± SE; 15,4 ± 2,6 cestodes). Fish mean wet-weight before and after parasites removale was respectively 17.27g (7.49-44.07 ± 1.61g) and 17.25g (7.48-44.03 ± 1.61g) with a mean difference before-after of 0.02g (0-0.055 ± 0.002g). Cestodes found directly in livers ranged from 0 to 36 (mean ± SE; 9,8 ± 1,7 cestodes). The populations’ liver cestode density differed significantly (*χ*^2^ = 17,223, df = 1, p-value = 3,324e-5; Fig. A1, available online).

Liver enzymatic activities of uninfected fish from Lake Triton (control) and Lake Cromwell (4 uninfected fish) did not significantly differ for CPT (p-value = 0,05395; Fig. 1A) and CCO activity (t-value = 2,0441, p-value = 0,06452; Fig. 1D). However, CS (p-value = 0,001998; Fig. 1B) and LDH (t-value = −5,8783, p-value = 0,004028; Fig. 1C) did differ between the two populations. Fish from Lake Cromwell showed a lower CS activity and a higher LDH activity than fish from Lake Triton.

**Figure 1.**
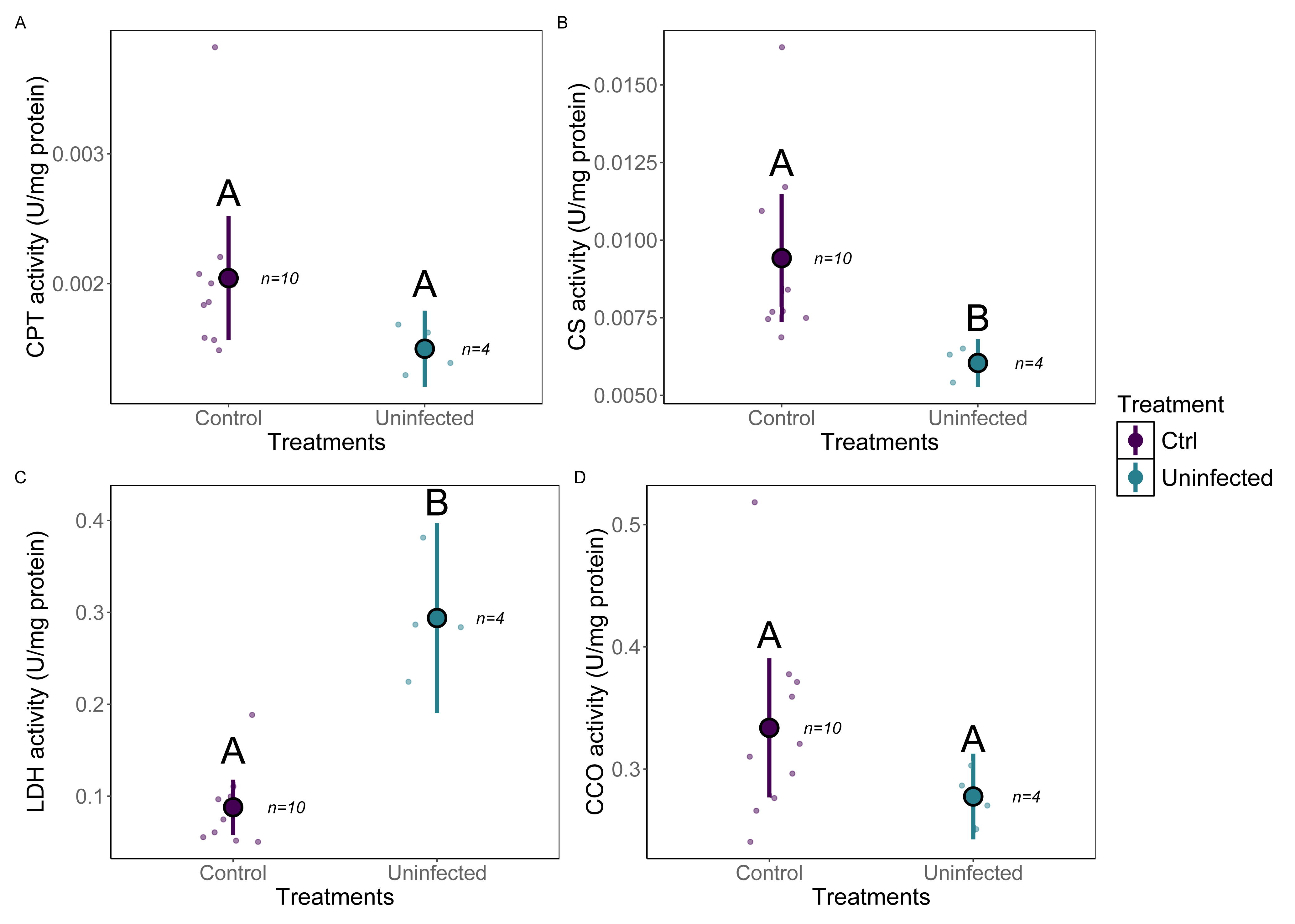
Liver enzymatic activities of uninfected fish from lakes Triton and Cromwell. Dots with black border represent mean values, lines represent 95% confidence intervals and colored dots represent actual data. Capital letters demonstrate significant differences between groups. A) Enzymatic activity of CPT, B) Enzymatic activity of CS, C) Enzymatic activity of LDH and D) Enzymatic activity of CCO.

When comparing fish liver enzymatic activities among treatment groups (i.e control, cleaned, repopulated, parasites only), we observed some significant differences between parasite and/or control livers and infected livers. CPT activity of liver tissues was similar across all groups (p-value = 0,2877; Fig. 2A). However, CPT activity of parasites was under the detection threshold of the spectrophotomer and similar to the negative control (reaction reagents without sample). Thus, comparisons for this enzyme were made only among the liver groups. CS activity varied significantly among groups (*χ*^2^ = 26,477, df = 3, p-value = 7,579e-6; Fig. 2B). Control fish livers had significantly higher CS activity than repopulated livers, but cleaned livers did not significantly differ from either control or repopulated livers (Table A2, available online). Parasite CS activity was significantly higher than hepatic tissues (Table A2, available online, and Fig. 2B).

**Figure 2.**
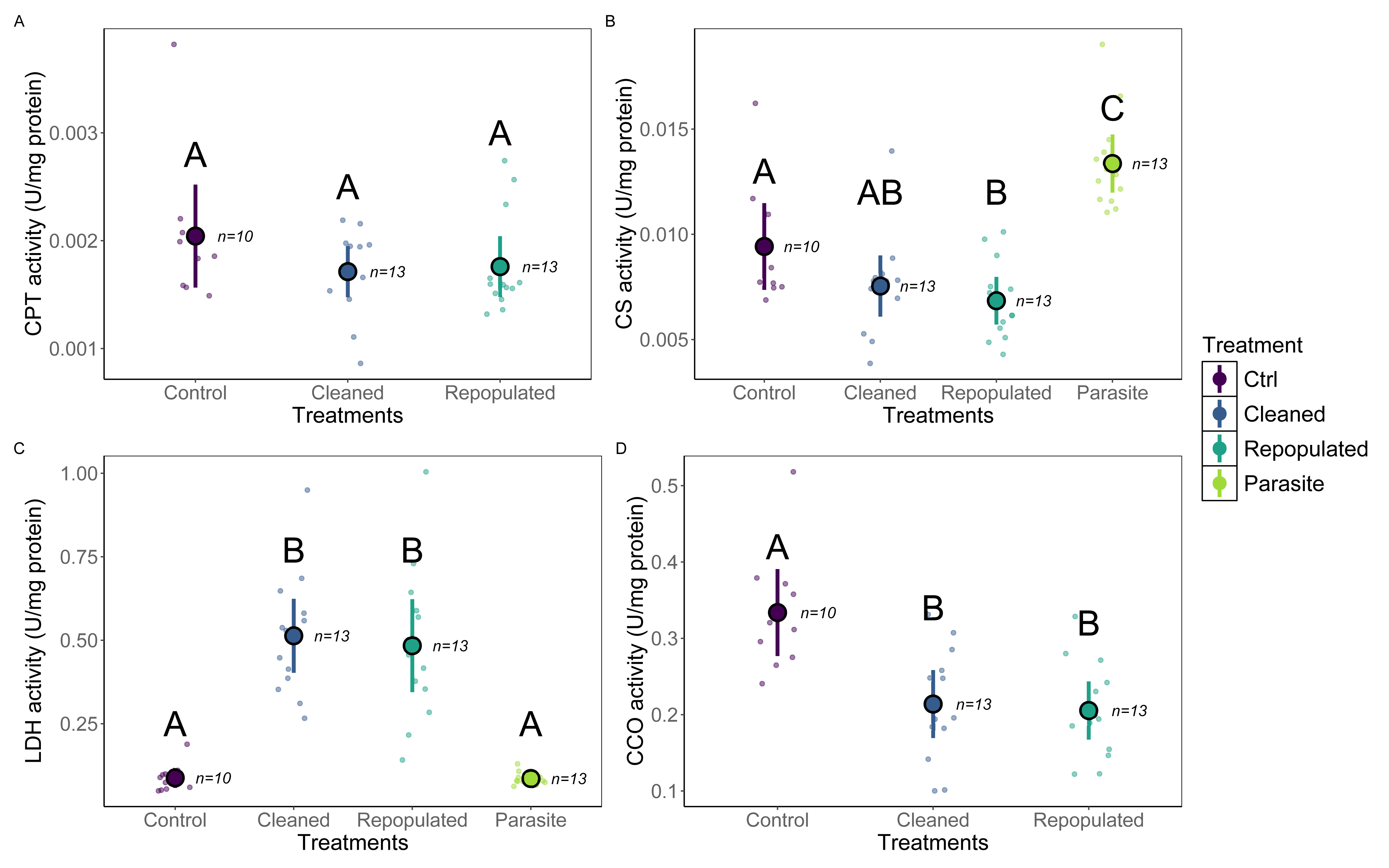
Enzymatic activities of control livers, cleaned livers, parasite-repopulated livers and parasites. Dots with black border represent mean values, lines represent 95% confidence intervals and colored dots represent actual data. Capital letters demonstrate significant differences between groups. A) Enzymatic activity of CPT, B) Enzymatic activity of CS, C) Enzymatic activity of LDH and D) Enzymatic activity of CCO. For CPT and CCO, there are only three treatments because parasites did not present CPT activity and due to a limited quantity of tissue CCO was not measured for parasites.

For LDH activity (*χ*^2^ = 35,713, df = 3, p-value = 8,61e-8; Fig. 2C), cleaned and repopulated livers had similar activities, but both were significantly higher than control livers (Table A2, available online). Parasite LDH activity was significantly lower than cleaned and repopulated livers, but similar to control livers (Table A2, available online, and Fig. 2C). For CCO activity, only the enzymatic activity of hepatic tissues was measured due to the lack of parasite tissue. Similar to LDH, CCO showed significant differences among groups (f-value = 9,843, df = 2, p-value = 0,000374; Fig. 2D): control livers had higher CCO activity than cleaned and repopulated livers and no difference was detected between cleaned and repopulated livers (Table A2, available online).

For all enzymes tested, there was no interaction between liver cestode density and treatment (Table A3, available online).The slopes had small non significant variations in the case of CPT (f-value = 0,088, p-value = 0,769; Fig. A4, available online), CS (f-value = 0,256, p-value = 0,618; Fig. A5, available online), CCO (f-value = 0,2857 p-value = 0,105; Fig. A6, available online), and LDH (f-value = 0,2927, p-value = 0,101; Fig. A7, available online).

The mitochondrial DNA of *Proteocephalus ambloplitis* (16S gene) was detected in all tested samples except for sample #3, which was from an uninfected fish, and served as a supplementary negative control. Every liver sample that was parasited showed a signal at ∼200 pb, as observed in the positive control (Fig. 3)

**Figure 3.**
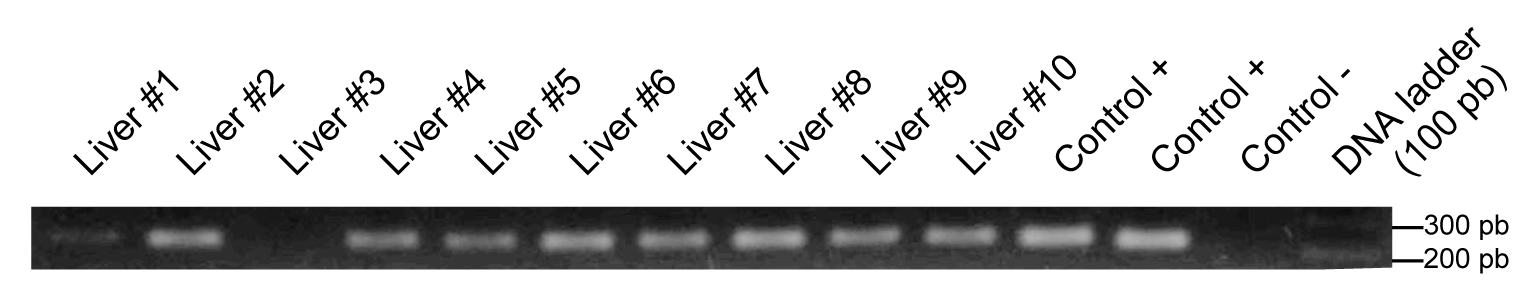
Evaluation of parasite contamination through PCR test in fish hepatic tissues. Parasite contamination was assessed via the amplification of the *Proteocephalus ambloplitis* mitochondrial 16S gene. Positive controls contain only parasite DNA and the negative control contained water. A molecular standard weight of 100 pb was used.

## Discussion

In response to the growing interest in studying cellular metabolism in response to parasite infections, our aim was to develop a protocol to purify parasite-infected livers to fully assess the impact of cestodes on hepatic enzyme activities. We studied three different groups of fish: (i) uninfected controls (fish from Lake Triton), (ii) cleaned infected fish (fish from Lake Cromwell for which parasites were removed from the liver), and (iii) repopulated infected fish (fish from lake Cromwell for which parasites were removed and put back with liver tissues) as well as the enzyme activity of the parasites in isolation. We also compared the activities between naturally uninfected fish from Lake Triton and Lake Cromwell to assess population-level differences in the tested enzymes unrelated to infection. Through these treatments, we aimed to test our protocol and evaluate if parasite contamination prevents the study of mitochondrial activity. We found that the activities of all enzymes tested did not differ between cleaned and repopulated livers, suggesting that contamination by parasite tissue is unlikely to alter the results of enzyme analyses in these tissues. However, in most cases, we observed differences in enzyme activities between uninfected (control tissues from Lake Triton) and infected fish (cleaned and repopulated liver tissues from Lake Cromwell) suggesting that infection may alter mitochondrial enzymatic activity in pumpkinseed sunfish. However, none of the enzymes were significantly correlated with infection load, and there were also significant differences in enzyme activities between naturally uninfected fish from lakes Cromwell and Triton, suggesting that population differences unrelated to infection exist. Lastly, DNA contamination was found in all livers that had previously contained parasites.

When comparing uninfected individuals from two different populations, differences in CS and LDH activity were observed. These variations could be explained by the presence of other parasite infections in 3 out of the 4 “uninfected” individuals. Even at low infection intensities (ranging between 0 and 54 parasites), enzymatic activities can be affected, particularly when compared to a population that has never been exposed to such infections (Mélançon et al. 2023; Sabbagh et al. 2024). The mere presence of parasites in the environment could account for these differences, as fish may allocate more energy toward immune responses (Rohlenová et al. 2011), leading to variations in enzyme activities. Moreover, uninfected individuals exposed to parasitism exhibit immunological responses, likely due to their hability to fight infections and succeding in staying uninfected (Sitjà-Bobadilla et al. 2008). This could explain the observed differences between our two populations and, in combination with our other results, suggests that parasite contamination is not the primary factor driving these population-level differences.

Parasite enzymatic activities were also analyzed to compare them with those of host liver tissues. Given that *P. ambloplitis* is an endoparasite, we expected it to mainly rely on anaerobic metabolic pathways due to limited oxygen availability in the liver. However, our enzymatic assays were not performed under anoxic conditions. When oxygen is available, cestodes can and will preferentially use their aerobic metabolism for energy production (Martínez-González et al. 2022). Indeed, we found elevated CS activity in parasites compared to host tissues, (Ryan et al. 2021)which suggests the utilization of aerobic pathways, such as the TCA cycle (Ryan et al. 2021). Given the mismatch between the test environment, where oxygen is present, and the host environment, which is typically anoxic, our results may not fully reflect physiological reality despite confirming that cestodes do have the capacity to metabolize substrates throguh the TCA cycle. Finally, as expected, no measurable CPT activity was detected in parasites, consistent with previous reports of CPT inactivity in cestodes (Barrett and Körting 1977). Indeed, cestodes are known to possess a non functional form of CPT (Barrett and Körting 1977), and its inclusion in our analysis was important for the validation of our protocol. If parasite tissue contamination in host liver samples had influenced our results, we would have expected to see differences between cleaned an repopulated livers. Parasite tissue could cause increased protein content or organ weight, which would have led to a decrease in normalized enzymatic activity. The normalization of enzymatic activity is a common practise in this field and we show that parasite contamination should not cause problems when implementing this practise.

We compared the enzyme activities of cestode parasites, uninfected livers, cleaned livers, and livers containing cestodes (repopulated livers). If parasites exhibited significantly lower enzymatic activity than their host’s hepatic tissues, we would expect their presence in liver samples to lead to an underestimation of the enzymatic activity of repopulated livers (i.e. essentially “diluting” the true fish activity). However, repopulated livers presented enzymatic activities similar to those of cleaned livers for CPT and LDH. Since cestodes alone had significantly lower activity than fish livers for these two enzymes, this suggests that parasite contamination did not lead to an underestimation of enzymatic activity in host tissues. Conversely, if parasites had higher enzymatic activities than hepatic tissues, we would expect an overestimation of enzymatic activity in repopulated livers compared to cleaned livers due to the presence of parasite tissue. However, for CS, where parasites had higher activity than fish tissues, cleaned and repopulated livers exhibited similar activities, indicating that parasite contamination did not lead to an overestimation of CS activity. Overall, our data show that the enzymatic activity of repopulated tissues do not differ from that of cleaned tissues and that parasite tissue metabolism does not interfere with host tissue activity in these assays. These findings support the reliability of previously published work (e.g. Sabbagh et al. 2024).

Based on prior research (Sabbagh et al. 2024), we expected to observe a mitochondrial response to parasite infections. Contrary to Sabbagh et al. (2024), no significant interaction between liver cestode density and treatment was observed for any of the tested enzymes (Fig. A6, available online), despite some non-significant trends between cleaned and repopulated livers for LDH, CS, and CCO activity. This decrepancy could be the result of a lower cestode abundance in our fish (median = 10,5) compared to the previous study (median = 26) (Sabbagh et al. 2024). The presence of cestodes in host organs is thought to cause tissular damage and to elicit immune responses, both of which could alter metabolic processes (Esch and Huffines 1973; Rohlenová et al. 2011; Deslyper et al. 2016; Mondal et al. 2016). Our results suggest that there could be a certain threshold below which parasite abundance is insufficient to produce detectable metabolic effects. Additionnaly, our relatively small sample size (n = 26) may have limited our hability to detect significant correlation. Nonetheless, the absence of a correlation between parasite abundance and enzymatic activities indicates that our protocol did not compromise liver tissue quality, since highly infected individuals exhibited similar enzymatic activity levels as those with lower infection levels. Although the time required to manipulate livers (and thus, the time between organ extraction from the fish and preservation in buffer) was greater the more infected they were, we found no evidence that these differences in manipulation time impacted the estimates of mitochondrial activity measured.

While there were no differences between cleaned and repopulated livers through enzymatic assays, PCR analysis revealed parasite contamination in all cleaned samples. Indeed, the mitochondrial 16S rRNA gene specific to *Proteocephalus ambloplitis* was detected in all samples except for one uninfected liver. This means that despite all our efforts to purify liver tissues, they remained contaminated with parasite DNA. Therefore, we cannot rule out the possibility that active mitochondria or cells from cestodes were present in cleaned liver samples. Regardless, the presence of parasite DNA does not seem to have impacted estimates of enzyme activities. These results are important to consider for futur projects that would require analyses of DNA taken from infected tissues.

Taken together, our results suggest that enzyme analyses on parasite-contaminated tissues can be conducted reliably, regardless of whether parasites are removed. However, this may not hold true for more highly infected samples. Indeed, the quantity of foreign protein is likely to be higher in heavily infected samples, potentially diluting host tissues sufficiently to bias enzymatic measurements. Since the parasite intensity threshold at which contamination begins to impact results remains unknown, we recommend the removal of parasite tissues from host tissue samples prior to assessing cellular metabolism activity to minimize possible biases.

## Conclusion

We developed a parasite purification protocol to evaluate the extent to which parasite contamination impacts estimates of enzyme activities in hepatic tissues of pumpkinseed sunfish infected with cestodes. Enzyme activities of purified liver tissues and contaminated livers were similar even if PCR analyses revealed presence of parasite DNA in purified livers. Our results suggest that the decontamination protocol was rigorous enough not to impact the accuracy of the enzyme activity estimates made. This protocol should be employed in future studies interested in understanding the effects of parasites on host cellular metabolism.

## Aknowledgments

We acknowledge the traditional lands of the Kanien’kehà:ka, Omàmiwinini, and Anishinabewaki First Nations on which the field and laboratory work for this project took place. We thank Gabriel Lanthier for logistic support at the SBL and his team. We thank Matthew Archambault, Thomas Gagnon, and Andréa Serres for their help during fish sampling. SAB and SB are supported by the Canada Research Chair program. SAB and SB are also supported through the NSERC Discovery Grants program. VM is supported by an NSERC PGS, an FRQNT and a Groupe de recherche interuniversitaire en limnologie (GRIL-PCR-22A07) research scholarships. TP was supported through an Undergraduate Research Award.

