## Appendix for "Evaluation of parasitic contamination and effects on estimates of mitochondrial enzymatic activities in infected fish livers"

**Appendix (available online)**

**Figure A1. Liver cestode density of fish from Lakes Cromwell and Triton.** Dots with black border represent mean values, lines represent 95% confidence intervals and colored dots represent actual data. Fish from Lake Triton had no cestodes in their liver. The liver cestode densities differed significantly between the two populations (p-value < 0,0001). Capital letters demonstrate significant differences between groups.
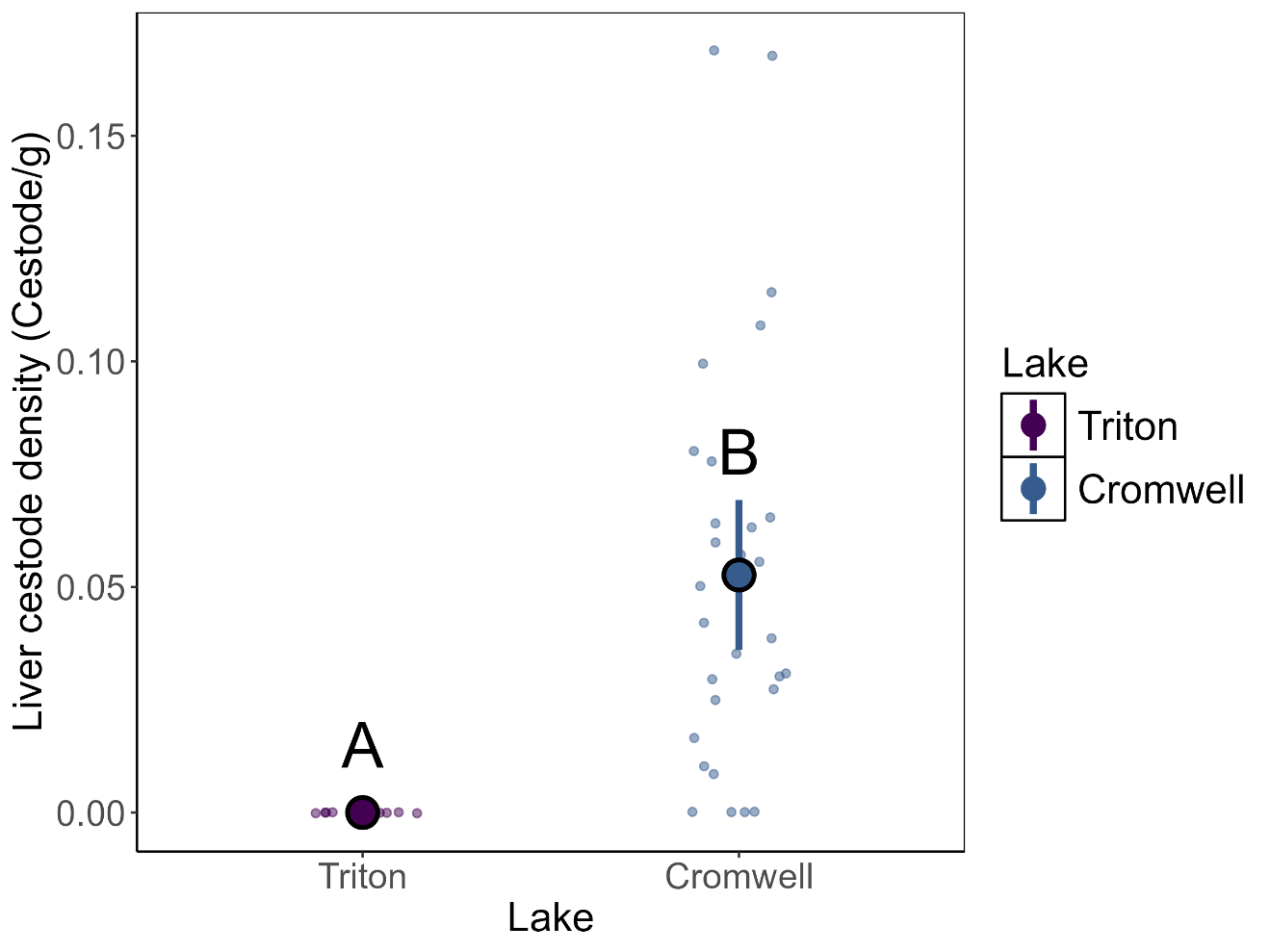

**Table A2. Kruskal-Wallis, Wilcoxon-Mann-Whitney, Anova and Tukey tests summary of liver enzyme activities, normalized on protein content, of all treatments.**

| Kruskal-Wallis test | | | | Wilcoxon Mann-Whitney | |
| --- | --- | --- | --- | --- | --- |
| Enzyme | Chi-Square | Degree of freedom | p-value | Treatment paired comparison | Holm’s corrected p-value |
| CS | 26,477 | 3 | 7,579e-06 | Ctrl-Cleaned | 0,37242 |
|  |  |  |  | Cleaned-Repopulated | 0,38969 |
|  |  |  |  | Cleaned-Parasite | 0,00013 |
|  |  |  |  | Repopulated-Ctrl | 0,03629 |
|  |  |  |  | Parasite-Ctrl | 0,00458 |
|  |  |  |  | Parasite-Repopulated | 1,2e-06 |
| CPT | 2,4918 | 2 | 0,2877 | Ctrl-Cleaned | 0,52 |
|  |  |  |  | Cleaned-Repopulated | 0,54 |
|  |  |  |  | Repopulated-Ctrl | 0,50 |
| LDH | 35,713 | 3 | 8,61e-08 | Ctrl-Cleaned | 7,0e-06 |
|  |  |  |  | Cleaned-Repopulated | 1 |
|  |  |  |  | Cleaned-Parasite | 1,2e-06 |
|  |  |  |  | Repopulated-Ctrl | 1,0e-05 |
|  |  |  |  | Parasite-Ctrl | 1 |
|  |  |  |  | Parasite-Repopulated | 1,2e-06 |
| ANOVA | | | | Test de Tukey | |
| Enzyme | F value | Degree of freedom | p-value | Treatment paired comparison | Ajusted p-value |
| CCO | 9,843 | 2 | 0,000374 | Ctrl-Cleaned | 0,0013711 |
|  |  |  |  | Cleaned-Repopulated | 0,9531430 |
|  |  |  |  | Repopulated-Ctrl | 0,0006377 |

**Table A3. General linear model of liver cestode density and treatment interaction for all enzyme activities of cleaned and parasite-repopulated livers of fish.**

| Enzyme | tte | Degree of freedom | F-value | P-value |
| --- | --- | --- | --- | --- |
| CPT | LCD | 1 | 0,614 | 0,442 |
|  | Treat | 1 | 0,015 | 0,905 |
|  | LCD : Treat | 1 | 0,088 | 0,769 |
| CS | LCD | 1 | 0,107 | 0,746 |
|  | Treat | 1 | 0,765 | 0,391 |
|  | LCD : Treat | 1 | 0,256 | 0,618 |
| LDH | LCD | 1 | 0,192 | 0,666 |
|  | Treat | 1 | 0,212 | 0,650 |
|  | LCD : Treat | 1 | 2,927 | 0,101 |
| CCO | LCD | 1 | 0,566 | 0,460 |
|  | Treat | 1 | 0,035 | 0,854 |
|  | LCD : Treat | 1 | 2,857 | 0,105 |

**Figure A4. Interaction between liver cestode density, treatment and carnithine palmitoyltransférase (CPT) activity of hepatic tissues.** CPT activity as a function of liver cestode density (number of cestode per gram of hepatic tissue). The model was not significant (F-statistic = 0,5271, p-value = 0,6676).
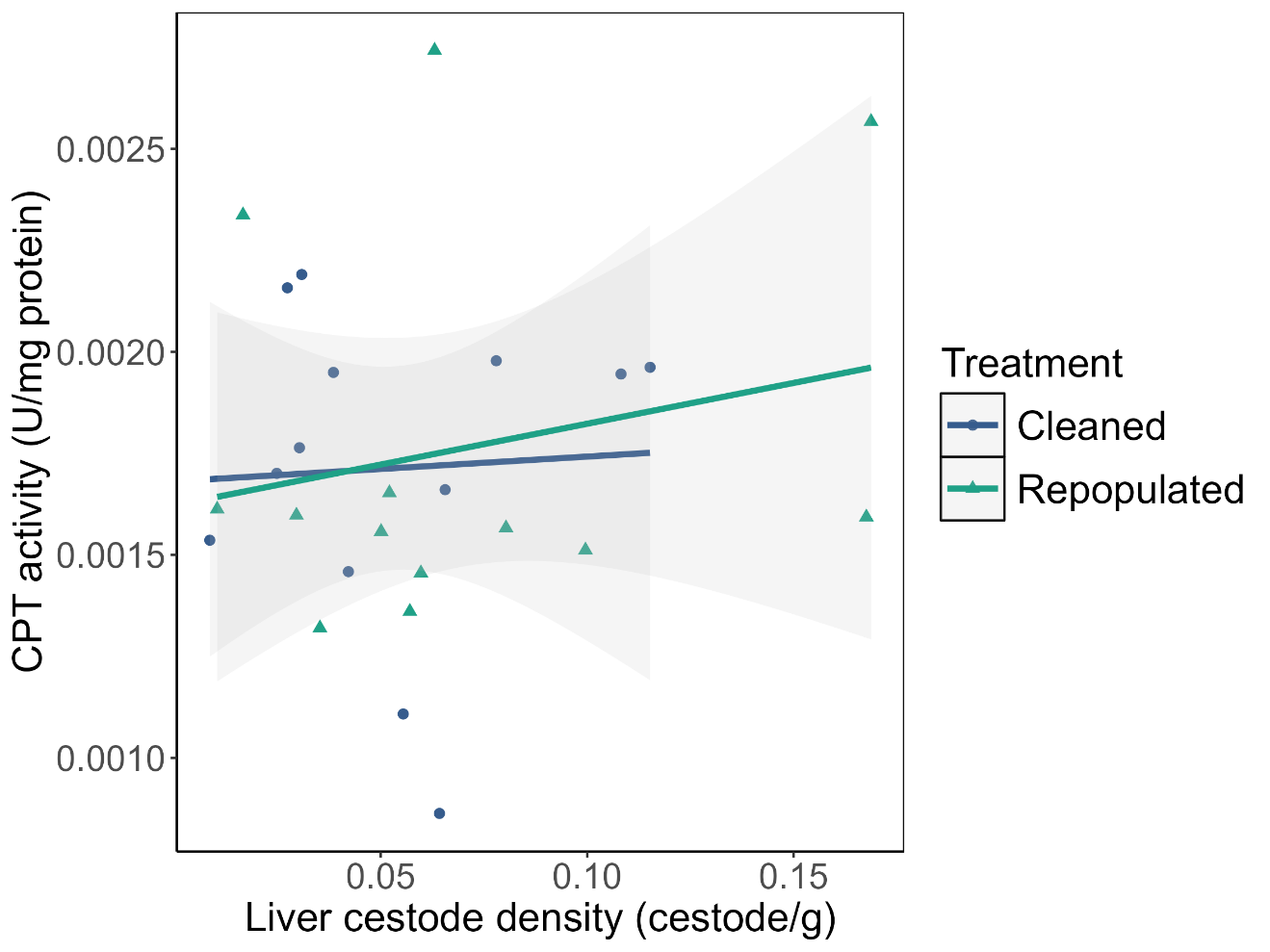

**Figure A5. Interaction between liver cestode density, treatment and citrate synthase (CS) activity of hepatic tissues.** CS activity as a function of liver cestode density (number of cestode per gram of hepatic tissue). The model was not significant (F-statistic = 0,6332, p-value = 0,6002).
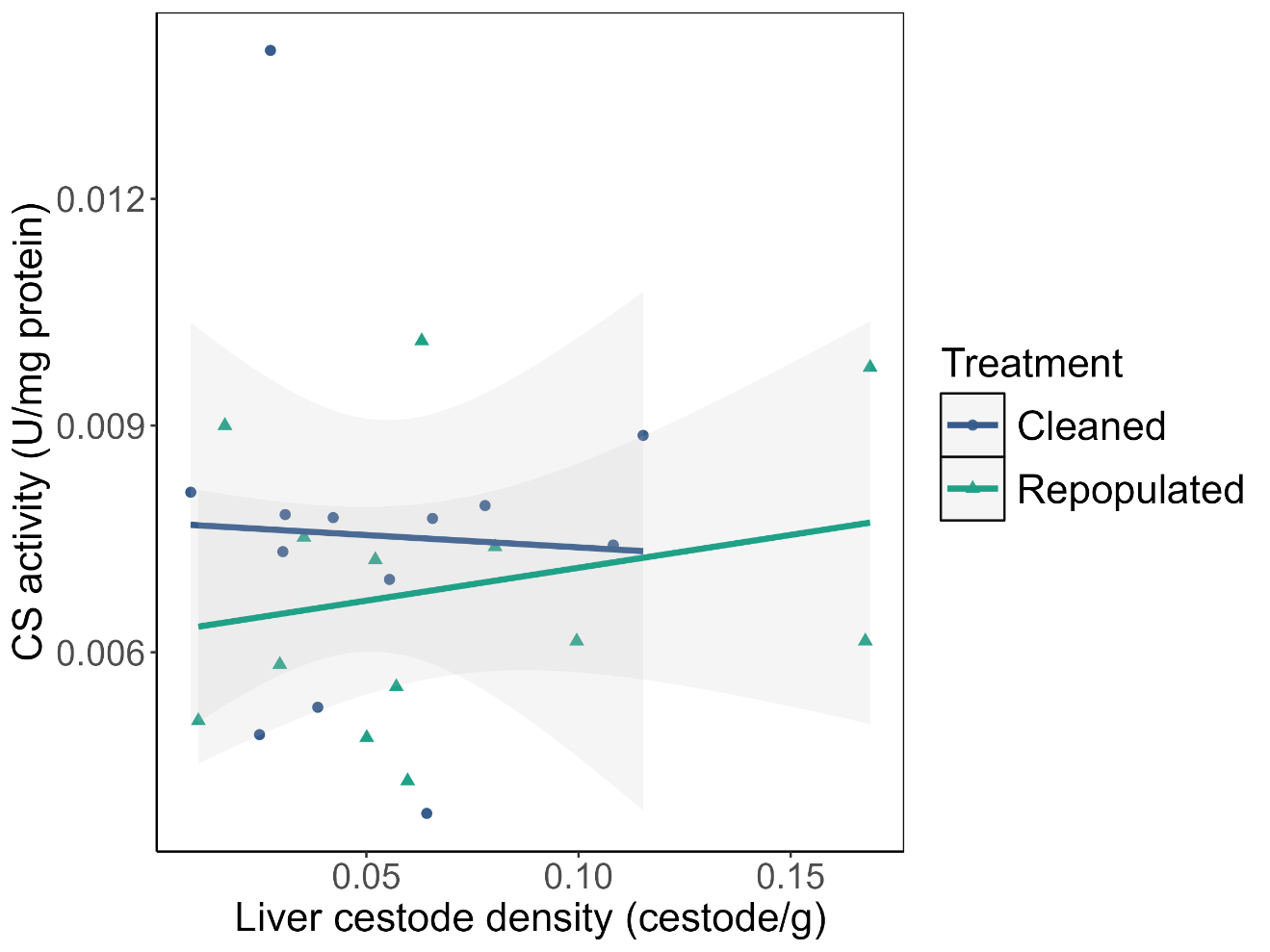

**Figure A6. Interaction between liver cestode density, treatment and cytochrome c oxidase (CCO) activity of hepatic tissues.** CCO activity as a function of liver cestode density (number of cestode per gram of hepatic tissue). The model was not statistically significant (F-statistic = 1,822, p-value = 0,168).
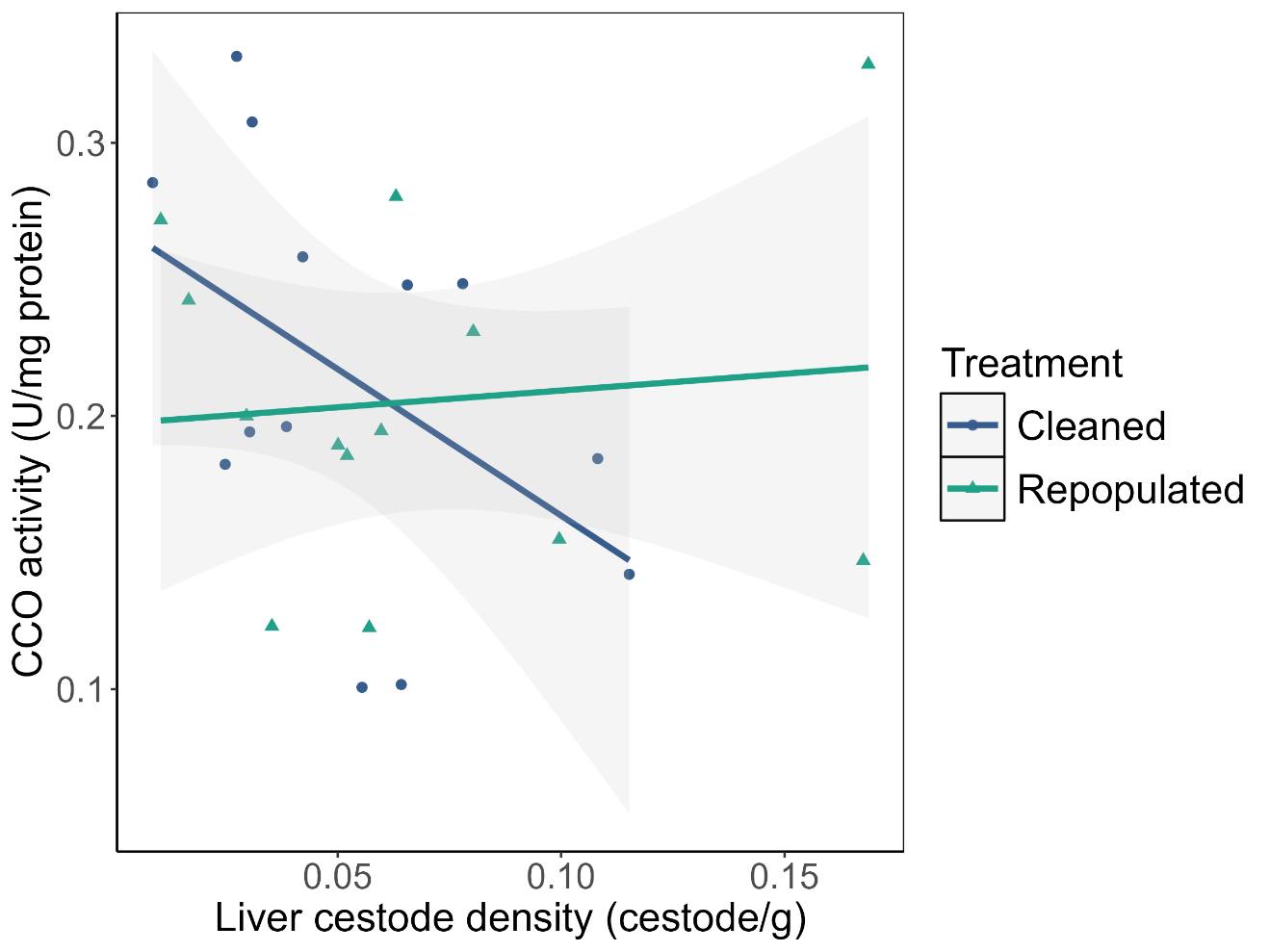

**Figure A7.** **Interaction between liver cestode density, treatment and lactate dehydrogenase (LDH) activity of hepatic tissues.** LDH activity as a function of liver cestode density (number of cestode per gram of hepatic tissue). The model was not significant (F-statistic = 1,63, p-value = 0,2067).
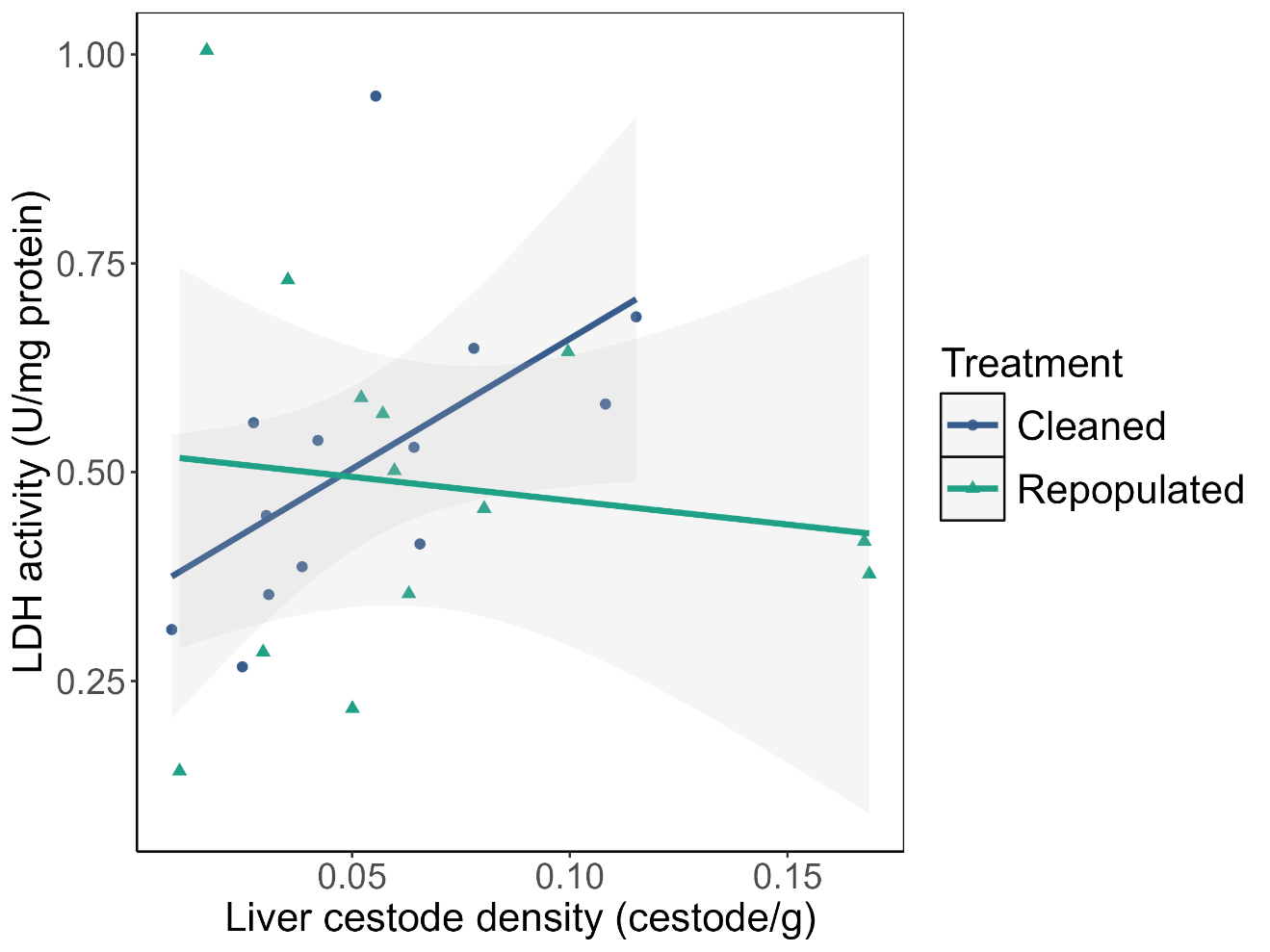
